# Asymmetric blood flow within a shared capillary network links the nucleus of the solitary tract and area postrema

**DOI:** 10.1101/2025.11.22.689934

**Authors:** Kaitlin E Carson, Ranjan Roy, Javier E Stern

## Abstract

The nucleus of the solitary tract (NTS) and the area postrema (AP) form a tightly coupled dorsal medullary complex that integrates visceral and humoral signals governing autonomic and cardiometabolic regulation. While their neural interconnections are well characterized, the organization and functional significance of their shared vascular network remain poorly understood. Here, we used *in vivo* two-photon microscopy in rats, combined with fluorescent vascular labeling and retrograde neuronal tracing, to visualize and quantify blood flow within the AP–NTS microcirculation. We identified direct capillary connections between the two regions, confirming a continuous vascular network previously inferred from ex vivo studies. Analysis of red blood cell trajectories revealed that blood flow across these junctions is predominantly unidirectional, from the NTS toward the AP. Morphometric measurements showed that AP capillaries are nearly twice the diameter of those in the NTS, implying a lower local vascular resistance and providing a structural basis for this directionality. Despite these geometric differences, capillary flow velocities were similar between regions, consistent with active regulation that maintains stable perfusion dynamics. Together, these findings uncover a previously unrecognized, functionally asymmetric vascular pathway within the dorsal medulla. By enabling the directed transfer of diffusible neurohumoral signals from the NTS to the AP, this specialized microvascular network adds a novel layer of communication between two key brainstem autonomic centers and may represent an additional mechanism for the integration of visceral and circulating information.

## Introduction

The nucleus of the tractus solitarius (NTS) and area postrema (AP) are two major brainstem visceral sensory and autonomic control nuclei that integrate and relay peripheral signals to hypothalamic and other forebrain regions that play a central role in the regulation of cardiometabolic functions^1–4^. Located in the dorsal surface of the medulla, the AP is a sensory circumventricular organ (CVO) that lacks a traditional blood-brain barrier, allowing for the direct sensing and processing of humoral signals^5,6^. In addition, the AP also borders the caudal wall of the 4^th^ ventricle, granting access to signals in the cerebrospinal fluid^7^. The NTS, located ventrolateral to the area postrema, is a large nucleus extending along the rostrocaudal length of the medulla and serves as the primary integrative center for vagal, glossopharyngeal, and spinal afferent inputs arising from multiple visceral organs.

The AP and NTS are not only topographically adjacent but also function as a coordinated dorsal medullary complex to integrate visceral and humoral signals governing cardiometabolic regulation. There are prominent neural connections linking the AP and the NTS. For example, circulating angiotensin II and vasopressin modulate glutamatergic neurons in the AP that project to medial NTS (mNTS) neurons, thereby influencing baroreflex-mediated sympathetic nerve activity^4,8,9^. In addition to direct neural connections, these regions share overlapping physiological functions. They are the primary central sites of action for glucagon-like peptide 1 (GLP-1) and amylin, whose receptors constitute molecular targets for pharmacotherapeutic interventions for the treatment of obesity and diabetes^10^. Recent studies have begun to elucidate the functional interplay between these two regions in feeding behavior, identifying their respective contributions to aversive and non-aversive mechanisms that reduce food intake^2,11^. This emerging work aims to inform strategies that minimize undesirable side effects of pharmacotherapies targeting obesity^12,13^. Thus, a deeper understanding of how these nuclei communicate with each other will facilitate the development of more effective interventions for complex physiological disorders such as obesity and cardiovascular disease.

A critical, though relatively underexplored, modality of communication between different brain regions is through diffusible signals. This is achieved in part through interconnected vascular networks, such as connected portal systems or capillary beds. In the brain, the first example of this is the shared portal system of the hypothalamus and the pituitary gland^14,15^, and more recent work has identified a secondary shared central portal system between the suprachiasmatic nucleus (SCN) and the organum vasculosum of the lamina terminalis (OVLT)^16,17^. These interconnected vascular networks are beneficial because they allow efficient transport of neurohormones and other peptides between capillary beds, bypassing the general circulation^15^.

Early studies characterized the anatomical vasculature irrigating the AP and NTS in rodents. Both the AP and the NTS receive their arterial supply from branches of the posterior inferior cerebellar artery (PICA), which then supply their respective capillary beds^18^. The AP contains a high density of capillaries that are larger than traditional brain capillaries^19^ due to fenestrations in the endothelial wall^5,6^ and double basement membranes^20^, characteristic of other CVO’s to allow for additional exposure to circulating factors^19^. These same enlarged capillaries are also connected to the surrounding choroid plexus^21^. Capillaries of the NTS, on the other hand, are comparatively smaller and unfenestrated^22^.

Notably, early work by Roth and Yamamoto (1968) proposed the existence of a portal venous system connecting the capillary beds of the AP and the NTS^18^. However, more recent detailed anatomical analyses by Yao et al. have demonstrated that, rather than a discrete portal system, capillaries from the AP extend directly into and merge with those of the NTS, forming an intertwined and continuous microvascular network^21^. This shared capillary network supports a new pathway by which diffusible signaling molecules could travel between the AP and NTS, at concentrations sufficient to elicit a physiological response in target neurons. Still, to fully elucidate the functional significance of the shared capillary network between the AP and the NTS, it is essential to determine the direction of blood flow within this network, which remains unknown. Establishing the flow dynamics within this shared capillary system will clarify whether a defined source/target relationship exists for diffusible signals transmitted through this vascular pathway.

To address this critical knowledge gap, we developed a novel *in vivo* rat approach to directly measure blood flow within the AP and NTS vascular systems. Using intravenous fluorescent dye labeling of the vasculature, combined with two-photon (2P) imaging, we examined both the anatomical organization and functional dynamics of the AP-NTS microvasculature in vivo. We quantified the rate and direction of flow within the arterioles, venules, and capillaries associated with each region. Importantly, we were able to visualize direct vascular connections between the enlarged AP capillaries and adjacent NTS capillaries, further supporting the presence of a shared capillary network linking the two regions. Finally, we found that blood flow within this network predominantly moves from the NTS towards the AP. Together, these studies demonstrate the presence of a functional shared capillary network, through which diffusible signals generated in the NTS can efficiently reach and influence the AP.

## Methods

### Animals

Male and female Sprague-Dawley rats (250-450 grams) were used for all experiments (n=24). Rats were housed in cages (2 per cage) under constant temperature (22 ± 2°C) and humidity (55 ± 5%) on a 12-h light cycle (lights on: 08:00-20:00). All performed experiments were approved by the Georgia State University Institutional Animal Care and Use Committee (IACUC) and carried out in agreement with the IACUC guidelines. At all times, animals had *ad libitum* access to food and water and all efforts were made to minimize suffering and the numbers of animals used for the study.

### Retrograde labeling of hypothalamic-projecting NTS neurons

Labeling of hypothalamic-projecting NTS neurons was achieved through microinjections of Lumafluor, a retrograde tracer, into the supraoptic (SON) and paraventricular (PVN) nuclei of the hypothalamus. Briefly, rats were anesthetized with intraperitoneal injection of a ketamine/xylazine (10%) cocktail (60 mg/kg) and monitored for cessation of the hindlimb withdrawal reflex. Body temperature was maintained using a heating pad. The top of the head was shaved, and the rat was fitted to a stereotaxic frame (Neurostar). The surgical area was prepped with iodine (1%) and ethanol (70%) scrubs and eye lubricant applied. An incision was made on the skin overlying the skull, and the skin was held back using bull clips, exposing the bregma suture line and lambda. Utilizing the Neurostar software to correct for tilt and scaling (ensuring scaling value is greater than 0.9), the left SON (one injection) and PVN (two injections) were localized using the following coordinates (anterior/posterior, medial/lateral, and dorsal/ventral) from bregma; -1.08, -1.83, and 9.30 (SON), -1.08, -0.28, and 7.16 (PVN #1), and -1.20, -0.23, and 7.70 (PVN #2). A hole was drilled in the skull over the injection sites (Carbide Burs, RA-7). Lumafluor (Green Retrobeads) was loaded into a 5 microliter Hamilton syringe with a 26-gauge, beveled needle and was placed using the manual micromanipulator (Neurostar). 0.4 microliters were then injected over a period of 7 minutes and waited at least 5 minutes post-injection before slowly removing the syringe needle. Once complete, the skin over the surgical site was closed using a combination of discontinuous suture (5-0 surgical suture) and glue (Vetbond). Animals were given a subcutaneous injection of carprofen for post-surgical pain (EthiqaXR, 0.5mg/kg) and body weight was monitored. The rats were left to recover at least one week before imaging experiments, this ensured the Lumafluor had time to travel and express in the NTS.

### In vivo two-photon (2P) imaging of the vasculature of the dorsal medulla

Rats were anesthetized with an intraperitoneal injection of urethane (1.5g/kg) and monitored for cessation of hindlimb reflex. Then the trachea was intubated, and the left femoral vein catheterized as described previously^23^. The rat was then fitted to a stereotaxic frame (Stoelting 51600 Lab Standard Stereotaxic Instrument). The dorsal brainstem was exposed via blunt dissection and cauterization (Thermal Cautery Unit, Geiger Medical Technologies), and a window created through removal of the meningeal membranes, ensuring calamus scriptorius was visible, modified from a previous approach to expose the vagal trigone^24^. To create space for the microscope objective, much of the occipital bone was removed using 1mm rongeurs, and a combination of superglue and hemostatic sponges (Vetspon) were used to move the cerebellum off the surface of the brainstem while keeping it preserved. The exposed brainstem was covered with optical gel and imaged under a 2P microscope system (Bruker Ultima Investigator, Bruker, Billerica, MA) excited with a Ti:Sapphire laser (Chameleon Ultra II (Coherent, Santa Clara, CA, USA)) tuned at 840 or 940 nm and scanned with resonant galvanometers through a 16X (numerical aperture 0.8) water immersion objective (Olympus, Center Valley, PA, USA). The medullary vasculature was labeled through intravenous infusions of either Rhodamine (20mg/mL) or fluorescein 70kDa dextrans (FITC, 20mg/ml). The AP was identified based on its characteristic triangular shape and the presence of large capillaries on the dorsomedial surface of the brainstem. The intermediate NTS was defined as the region lateral and ventral to the AP and would include Lumafluor-positive cells and smaller capillaries (see ***Fig.2***)

### Infusion of sodium fluorescein

To further confirm the location of area postrema, intravenous infusion of sodium fluorescein (NaF) was performed as described previously^25^. Briefly, NaF (2.5mg/ml), a small molecule dye that has been demonstrated previously as permeable to the fenestrated capillaries of area postrema^25^, was infused via a femoral vein catheter. After 90 minutes, to allow time for the NaF dye to begin infusing outside of the capillaries, imaging of the brainstem through a 4X (numerical aperture of 0.13) air objective was performed.

### Identification of venules, arterioles, and capillaries

The artery-specific dye Alexa Fluor 633 (AF 633) was used, as described previously^23,26^ to label arterioles in the AP and NTS. In a subset of rats, Alex Fluor 633 (1mg/kg) was intravenously injected at least 90 minutes before imaging to allow for labeling of elastin fibers that densely line arterioles. Once imaging, FITC dextran (20 mg/ml, 200 μl/rat) was intravenously injected to visualize all vasculature, and arterioles were identified with labelled elastin fibers. Those without AF 633 labeling were considered venules or capillaries based on their size and location.

### Measuring vasculature diameter, RBC velocity, and direction of flow

Images and videos captured to measure vasculature diameter, RBC velocity, and direction of blood flow were done using the Prairieview software (Bruker, Billerica, MA) in the Galvanometric scanning mode. To get baseline measurements of vessel diameter, z-stack images were acquired (512×512 pixels area imaged). Then, using Fiji software, lines were drawn through the width of the vessel to measure its diameter. Arteriole diameters were pooled into 2 main groups: large arterioles (50-100 μm, arising from lateral aspects of the medulla) and branching arterioles (20-50 μm, arising off the large arterioles). The central and lateral veins were defined based on their anatomical location. Red blood cells (RBCs) appear as dark spots in the fluorescent dye-filled vessels. To measure the direction these cells are flowing, as well as their velocity, line scans were performed. Briefly, a freehand line (30-100μm) was drawn running parallel to the vessel walls at 2X magnification and scanned (line scan duration of ∼2s and a sampling frequency of 5 kHz). The kymograph generated from the line scan, where RBC’s can be identified as the black stipes, was uploaded into Fiji software for further analysis. The direction of flow was determined by looking at the polarity (positive or negative) of the slope of the RBC’s striations through the kymograph. To calculate the velocity of the RBCs, they were first identified as dark bands on the kymograph. Then, a line was manually draw using the line tool to generate the angle of the line (the angle between the drawn line and horizontal axis), taking care to always draw the line in the same direction. This angle was then used to calculate the angle directly opposite from the x axis (i.e distance). After converting this angle to radians, now represented as Ill, the following equation was used to calculate velocity:

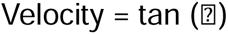

Once the velocity ratio was calculated, it was converted into μm/ms using the embedded distance (μm/pixel) and line period (ms/pixel) values from the line scan. This same procedure was done 20 times for each kymograph, sampling a different band each time to account for RBC velocity variability within the time sampled. Then the velocity values were averaged for every line scan at each time point.

### Fluorescence immunohistochemistry

A subset of rats (n=2) that received the Lumafluor microinjections was used to confirm expression of Lumafluor fluorescence in medial NTS neurons. Rats were anesthetized with an i.p injection of pentobarbital (Euthasol, 80mg/kg) and perfused transcardially with 0.1M PBS followed by 4% paraformaldehyde. For fluorescence immunohistochemistry, the medulla was dissected and blocked (O.C.T., Fisher Healthcare), following which 40 micron-thick serial coronal sections were collected using a Leica Cryostat and stored in 0.1M PBS until use.

A standard immunohistochemistry protocol was used to label neurons that contain tyrosine hydroxylase (TH), a marker for catecholaminergic neurons and a known subtype of hypothalamic-projecting NTS neurons^27^. Briefly, free floating sections were washed in PBS (0.1M) then incubated in a buffer (0.1% Triton X-100) and normal donkey serum (5%) for 1 hour. Slices were then incubated in an anti-TH monoclonal primary antibody (1:2000; Sigma-Aldrich) for 3 days at room temperature. Sections were then incubated in Alexa Fluor 594-conjugated donkey anti-mouse secondary (1:500; Jackson ImmunoResearch) overnight and mounted on subbed slides for imaging.

### Confocal imaging of fixed tissue

Images were captured using a laser scanning confocal microscope (Zeiss LSM 980 with Airyscan 2). Z-stacks of coronal sections of the medulla that contained the intermediate NTS (defined as the NTS present with the AP) were captured at 20X magnification, using the tiling feature to cover the NTS region. An average of 10 optical sections were collected per z-stack (with 3μm steps), and each animal had on average 3 coronal slices that would contain the intermediate NTS. During image acquisition, the master gain and laser power were kept consistent among samples. Quantification of Lumafluor+/TH+ and Lumafluor+/TH- cells in the intermediate NTS were performed in Fiji software, with cells with fluorescence signal from both considered co-localized.

### Statistics

Statistical analyses were carried out using GraphPad Prism 9 (GraphPad Software, California, USA). To compare groups, various tests were employed, including Student’s unpaired t-test and one-way analysis of variance (ANOVA) followed by Tukey post-hoc tests for multiple comparisons. The results are denoted as mean ± standard error of the mean (SEM). Statistical significance was established at p < 0.05.

## Results

### Implementation of a new approach to visualize in vivo the vasculature irrigating the AP and medial NTS

Anatomical maps of the vasculature supplying the AP and NTS regions have previously been created in a mouse model^21^. The direction and rate of blood flow, however, have not been characterized in this region. To accomplish this, we implemented a new experimental rat model that enabled us to expose the dorsal brainstem region in an anesthetized rat. In this preparation (***Fig.1A***), we then used in vivo two-photon (2P) imaging to anatomically and functionally assess the vasculature of the AP and NTS region, to determine blood flow directionality and rate of the major arterioles and venules found in these regions. To this end, the brainstem was exposed, and the calamus scriptorius (the space at which the central canal and the 4^th^ ventricle meet) was used as a centering point to find the caudal aspect of the AP. This region is also readily recognizable by the grey coloration of the tissue through a brightfield microscope (see ***Fig.1B, upper panel***). Vessels were labeled with fluorescent markers (Rhodamine-70 or FITC-70, 0.2 ml, i.v., ***Fig.1B, lower panel***). Under 2P imaging, the AP was readily identifiable by a triangular shape defined by the dense bed of large, fenestrated capillaries near the dorsal surface of the brainstem (***Fig.1B*,*2A***).

**Figure 1:**
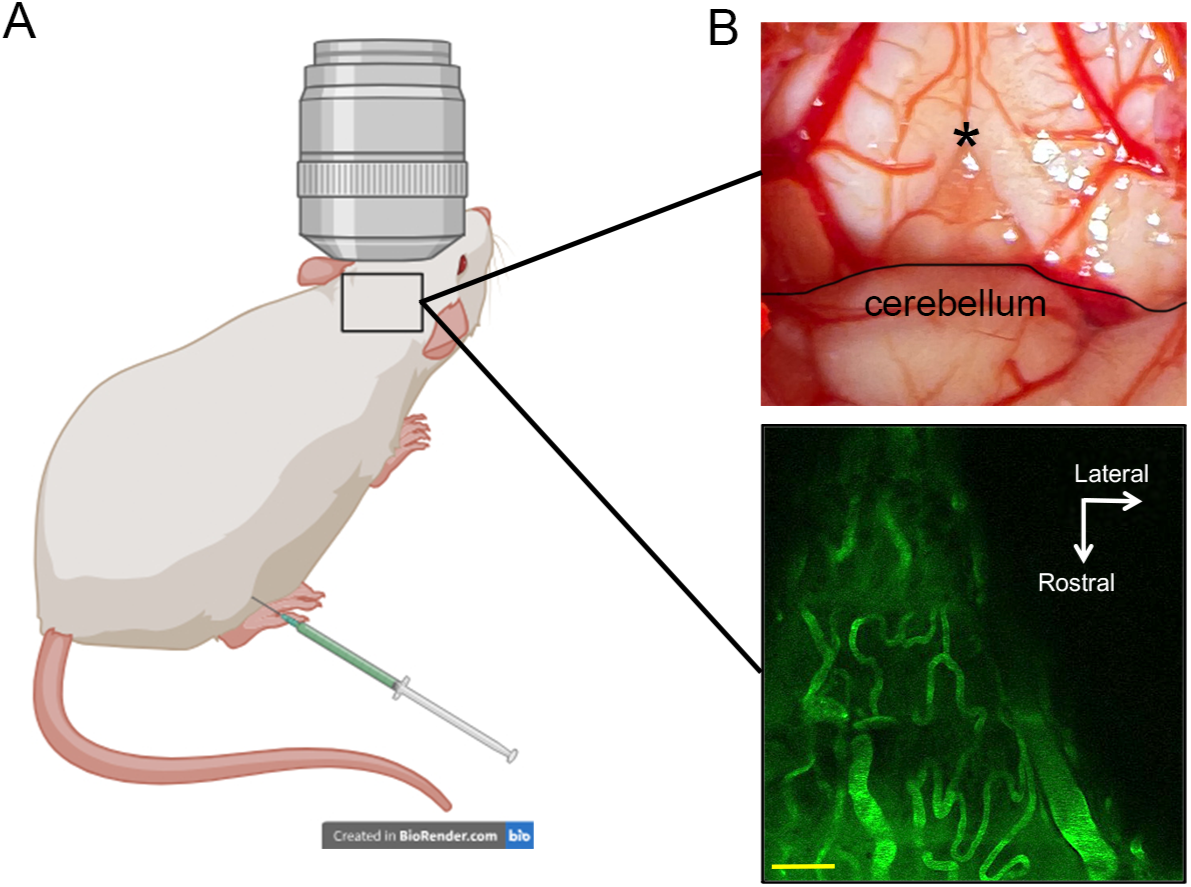
Experimental preparation for two-photon (2P) microscopy imaging of the dorsal brainstem. **A**. Graphic showing the experimental preparation set-up, including placement of the 2P objective above the brainstem in an anesthetized rat with combined femoral intravenous injection of fluorescent vessel dyes. **B**. Top: representative image of the exposed dorsal brainstem after microdissection. * denotes calamus scriptorius, the caudal edge of area postrema (AP). The image shows an intact cerebellum. Bottom: representative image of intravenous-dye-filled blood vessels within the AP. Scale bar= 50 μm.

**Figure 2:**
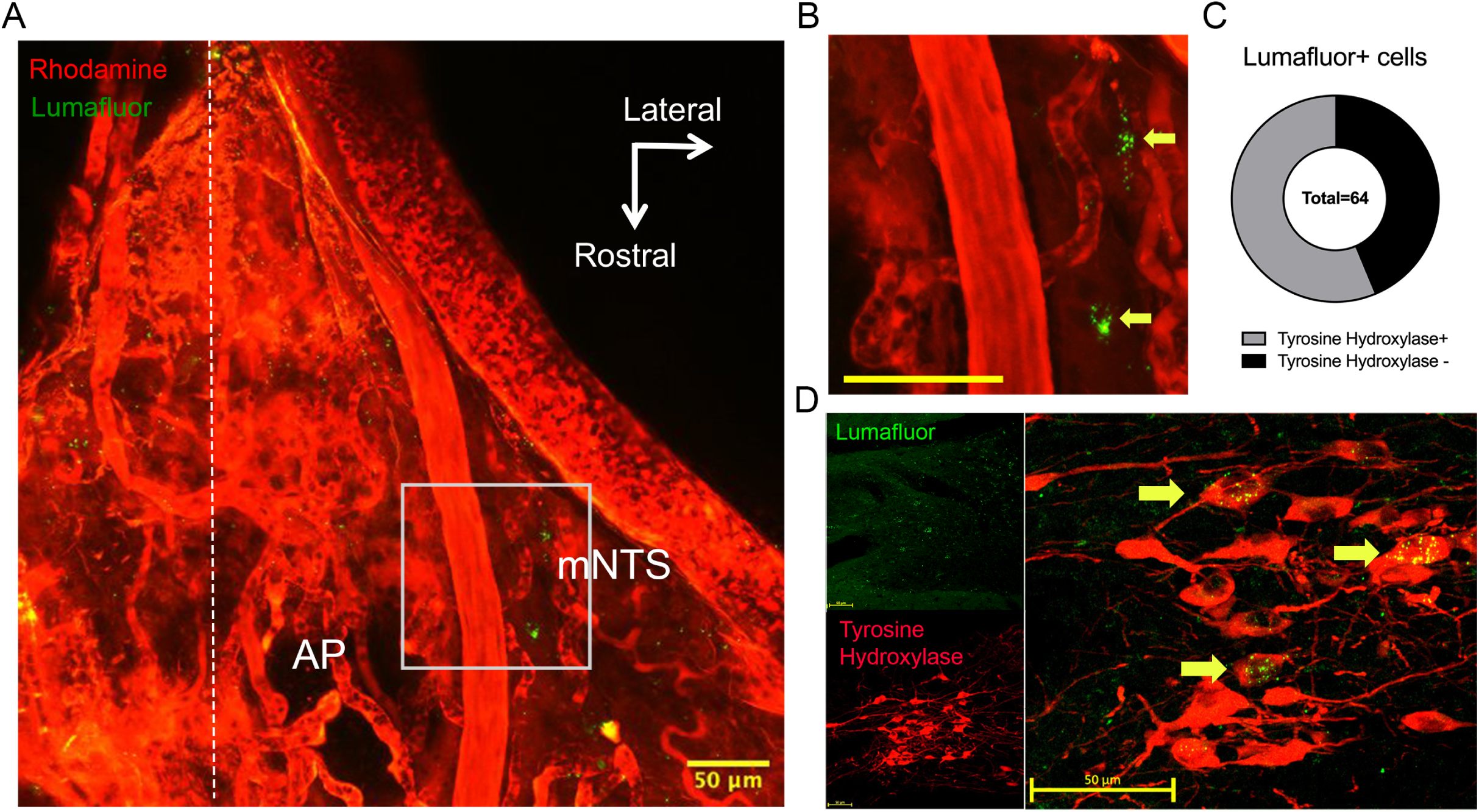
mNTS neurons identified in the vicinity of the labeled vasculature are primarily catecholaminergic. **A**. Representative image of the rhodamine-labelled (red) vasculature in the AP and mNTS region, along with retrogradely-labeled (green) NTS neurons present within the call-out box. Midline denoted by dashed white line. **B**. Call-out box from (A) with retrogradely-labeled (green) NTS neuron pointed out with yellow arrows. Scale bar= 50 μm. **C**. Of the 64 retrogradely-labeled NTS neurons counted, 36 (56.25%, grey) were immunoreactive to tyrosine hydroxylase. The remaining 28 (43.75%, black) were not. **D**. Representative images of medullary coronal sections showing retrogradely-labeled neurons (green) and tyrosine hydroxylase-labeled neurons (red). Co-localization of the signal denoted by yellow arrows.

To determine where the medial NTS (mNTS) lies in relation to the AP in this preparation, a subpopulation of hypothalamic-projecting NTS neurons was retrogradely labeled via injections of Lumafluor unilaterally into the supraoptic (SON) and paraventricular (PVN) nuclei (n=6 rats). Hypothalamic-projecting NTS neurons were readily identifiable under the 2P and were characterized by the fluorescent puncta congregated within a cell. In all cases, they were located lateral and ventral to the AP capillaries (***Fig.2A,B***). A combined retrograde tracing and immunohistochemistry for tyrosine hydroxylase study showed that 56.25% of the mNTS neurons labeled were catecholaminergic (***Fig.2D***), which aligns with previous reports stating that 40-70% of hypothalamic-projecting NTS neurons are catecholaminergic^28,29^.

Additionally, further confirmation to differentiate the AP from the mNTS was achieved through the use of intravenous NaF, a fluorescent dye that has been previously shown to be permeable to the capillaries of AP and not the NTS^25^. Supporting these previous observations^25^, the NaF dye excised from the regions containing the enlarged capillaries, further confirmation of the location of AP (***Supplementary Fig.1***). Taken together, the combined use of anatomical landmarks with retrograde tracing and permeable dye approaches supports our ability to identify and differentiate the AP and mNTS for subsequent functional vascular mapping.

### Identification and characterization of anatomical and functional properties of arterioles, venules, and capillaries in the AP and mNTS

To differentiate the various vessel types (i.e., arterioles, venules, and capillaries), we infused Alexa Fluor 633, which binds to elastin fibers to label arterioles^26^ as we previously used to successfully differentiate arterioles from venules in the SON^23^. Alexa Fluor 633 readily labeled two major arterioles originating from the posterior inferior cerebellar artery (PICA), which run from the lateral brainstem and head rostrally and parallel to the midline, on both sides (***Fig.3A***). These were found to be the largest arterioles present in the AP/NTS region, with an average diameter of 70.82 ± 2.99 μm (n= 6, from 5 rats) (***Fig.3D***). After performing a line scan of the arterioles and generating a kymograph to track RBC movement, we determined that blood flowed through these arterioles medially and rostrally (***Fig.3F***). These large arterioles gave rise to various smaller parenchymal arterioles with an average diameter of 28.35 ± 1.66 μm (n= 8, from 5 rats) that “dove” perpendicularly towards the midline, projecting along the same path as a midline-running venule (caudally, upstream to the AP, see ***Fig.3F***), directly giving rise to the AP capillary bed. Rostral arteriole branches, at the level of AP, similarly head towards midline before projecting ventrally to become continuous with NTS capillaries. As expected, the blood flow direction of all these smaller arteriole branches is from the lateral arterioles towards the capillary beds they supply (***Fig.3F***). These smaller parenchymal arterioles were found to have faster RBC velocities (18.80 ± 2.79 μm/ms n= 7, from 5 rats) compared to the lateral arterioles (8.19 ± 0.83 μm/ms n= 5, from 5 rats; p=0.0104, *post-hoc* Tukey’s multiple comparisons test; ***Fig.3E***).

**Figure 3:**
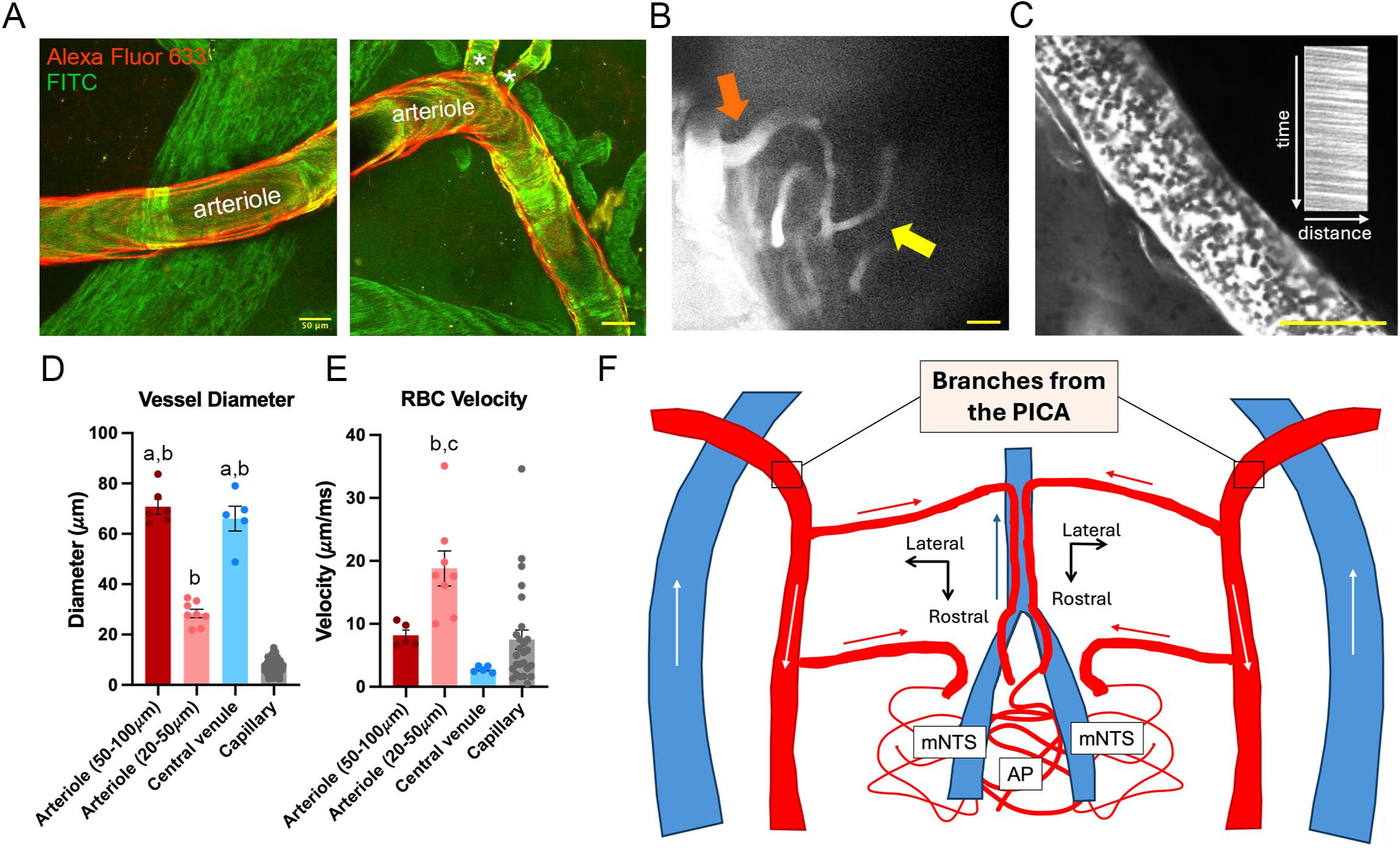
Functional vascular map of the AP and NTS regions. **A**. Representative composite of images displaying a large arteriole labeled with Alexa Fluor 633 (red) and i.v. FITC (green). Branching parenchymal arterioles are indicated with *. Scale bar= 50 μm. **B**. Representative image of an arteriole (orange arrow) directly supplying a smaller capillary (yellow arrow) within the mNTS. Scale bar= 20 μm. **C**. Red blood cells visualized within a vessel (black circles) shown with an example of a kymograph generated from the line scans of the blood vessel, showing striations that represent the RBCs moving across space (x axis) and time (y axis). The polarity of the slope of these RBC lines, in this example, a negative slope, represents the direction of RBC flow. Scale bar= 50 μm**. D**. Diameter measurements of the large lateral arterioles (dark red, n=6), small branching arterioles (light red, n=8), central venule (light blue, n=5), and capillaries (grey, n=164). The lateral arterioles and central venules are significantly larger than the small branching arterioles, and all have significantly larger diameters than the capillaries. (‘a’ denotes p < 0.05 compared to arteriole (20-50μm) and ‘b’ denotes p < 0.05 compared to capillary via one-way ANOVA followed by Tukeys post-hoc test for multiple comparisons)**. E**. RBC velocity measurements of the large lateral arterioles (dark red, n=5), small branching arterioles (light red, n=7), central venule (blue, n=5), and capillaries (grey, n=26). The small branching arterioles have a significantly faster RBC velocity than both the capillaries and central venule. (‘c’ denotes p < 0.05 compared to central venule via one-way ANOVA followed by Tukeys post-hoc test for multiple comparisons). **F.** Proposed microcirculation map of the AP and mNTS regions. Arterioles are represented in red, venules in blue. The direction of the blood flow is indicated by arrows within each vessel population. The path of the large lateral arterioles can be seen crossing over large veins on the lateral edge and continuing rostrally. From there, small branching arterioles can be seen heading towards the midline (dashed vertical line) and supplying both the AP and the NTS, with blood flow directed towards their capillary beds. The central venule is shown traveling caudally along the midline, with blood flow in the same direction.

We identified two major groups of clearly defined putative venules; a large venule located in lateral aspects of the brainstem, running parallel to the larger arterioles described above, and a group of smaller venules located medially, running at the border region between AP and NTS, and then following along the midline. Compared to the neighboring parenchymal arterioles, their mean diameter (66.01 ± 4.91 μm n= 5, from 5 rats) and mean RBC velocity (2.78 ± 0.22 μm/ms n= 5, from 5 rats) were significantly larger (p < 0.0001) and slower (p=0.003, *post-hoc* Tukey’s multiple comparisons test; ***Fig.3E***) In all venules, blood flowed in the caudal direction (i.e., opposite to the arterioles), allowing these different vessel types to be also differentiated based on their blood flow directionality (***Fig.3F***) in the absence of Alexa Fluor 633 labeling. A cartoon vascular map summarizing these findings is shown in ***Fig.3F***.

As shown in Figures 1B and 2A, the mesh of capillaries in the AP/NTS region was readily identified following intravascular labeling with rhodamine or FITC. Capillaries in the AP/NTS region had a mean diameter of 7.31 ± 0.21μm (n= 164, from 5 rats) and a mean RBC velocity of 7.51 ± 1.52μm/ms (n= 26, from 5 rats), significantly smaller (p<0.0001, ***Fig.3D***) and slower than the small parenchymal arterioles (p=0.0015; *post-hoc* Tukey’s multiple comparisons test; ***Fig.3E***).

### The AP and mNTS capillary beds are interconnected, with blood flowing predominantly from the mNTS towards the AP

In a recent, detailed anatomical study using mice, Yao et al. provided conclusive evidence for interconnected capillary beds between the AP and mNTS^21^. Still, whether this is also the case in rats, and more importantly, whether blood flow between these beds follows a particular direction, remains unknown.

Using the new experimental preparation described in this paper, we aimed to address these gaps. Similar to what was described anatomically in the mouse^21^, we confirmed in that in the in vivo rat brainstem, AP capillaries have a significantly larger diameter (8.95 ± 0.21μm n=96, from 5 rats) than NTS capillaries (4.99 ± 0.17μm; n=68, from 5 rats p<0.001, t(162)=13.93, two-tailed unpaired t-test; ***Fig.4A-C***). Conversely, and despite these differences in diameter, there were no differences observed in the RBC velocity between AP (8.32 ± 2.712 μm/ms, n=13 from 5 rats) and NTS capillaries (6.69 ± 1.469 μm/ms; p=0.6035, unpaired t-test; ***Fig.4D***).

**Figure 4:**
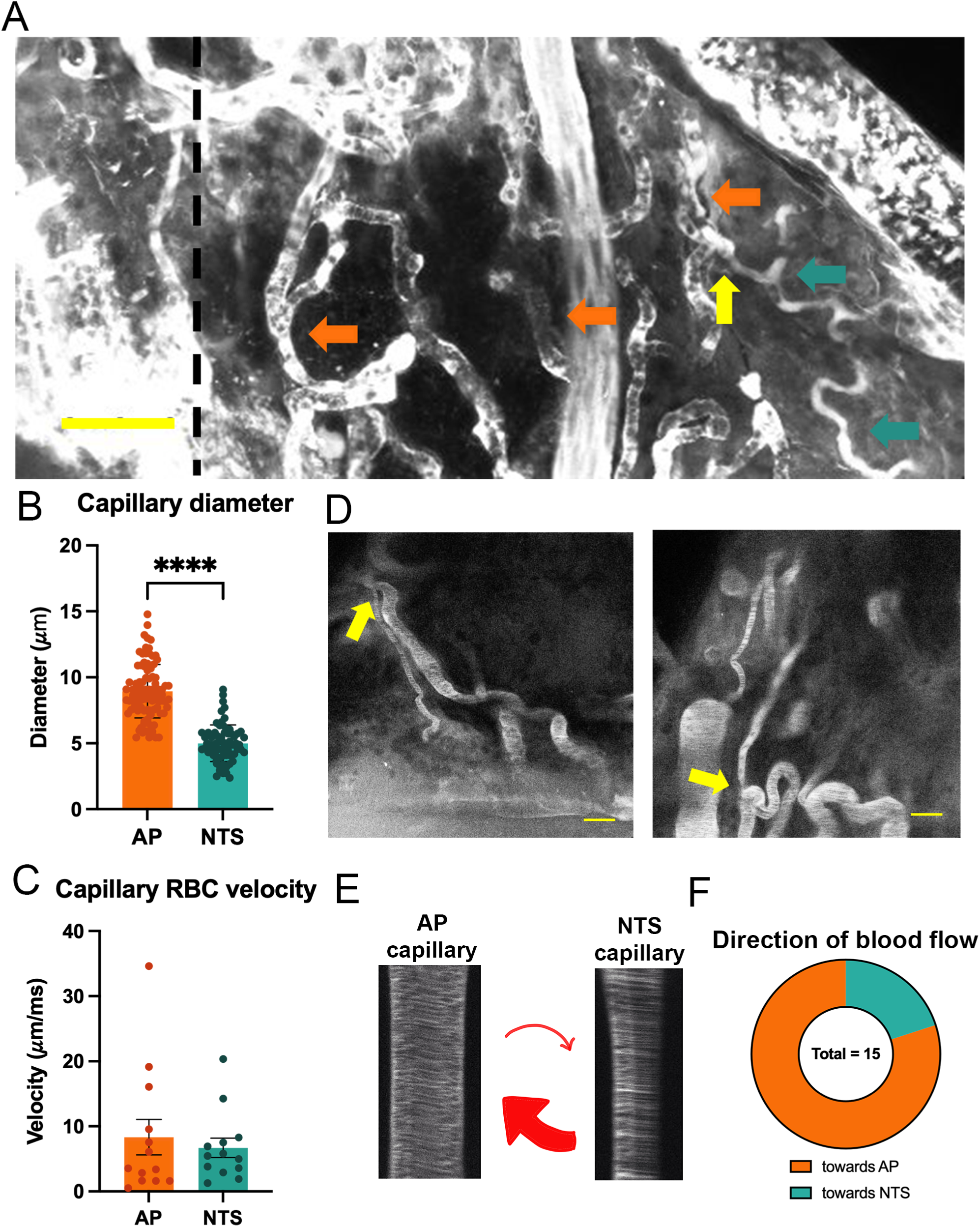
Blood flows predominantly from the NTS to the AP in their connected capillary beds. **A**. Representative image highlighting capillaries within the AP (orange arrows) and mNTS (teal arrows) at the border of these two regions. The dashed vertical line notes the midline. A junction point is shown with the yellow arrow, where the connected capillary clearly decreases in size as it transitions from the AP to the mNTS. Yellow scale bar= 50 μm. **B**. Diameter measurements of AP (orange, n=96, 5 rats) and mNTS (teal, n=68, 5 rats) capillaries. AP capillaries are significantly larger than mNTS capillaries. **** p < 0.001 via Students t-test. **C**. RBC velocity measurements of AP (orange, n=13, 5 rats) and mNTS (teal, n=13, 5 rats) capillaries. The RBC velocity of these two capillary populations is not significantly different (p= 0.6035, via unpaired t-test). **D**. Representative images of connected AP (larger) and mNTS (smaller) capillaries, pointed out with yellow arrows. Capillaries are filled with FITC for subsequent line scanning. Scale bar= 20 μm. **E**. Representative kymographs from line scans determining the direction of blood flow from a connected AP and NTS capillary. Red arrows indicate predominance of flow from NTS to AP. **F**. Fifteen connection points between AP and mNTS capillaries were identified in 3 rats and line scans performed on both sides of the connecting point to determine the direction of blood flow. A predominant (n=12, 80%) of those connection points measured showed blood flowing from NTS to AP (orange), with a smaller proportion (n=3, 20%) flowing from AP to NTS, demonstrating a non-unilateral but preferential flow from NTS to AP.

To establish the direction of flow between the two capillary beds, we aimed to identify precise junction points between a large (AP) and a small (NTS) capillary. Using this approach, we identified 15 junction points across 3 rats. While it was highly challenging to find them, once we did, they were clearly identifiable and located in the border area between these two regions. Some examples of these AP-NTS connected capillaries are shown in ***Fig. 4A and B***. Ultrafast line scans were taken on both sides of the capillary junction, and kymographs were generated to determine blood flow directionality (***Fig.4E***). Our results indicate that in the vast majority of the junctions (12/15 from 3 rats, 80%), blood flowed from the NTS side towards the AP side (***Fig.4F*)**.

## Discussion

In this study, we used a combination of 2-photon microscopy and infusion of fluorescent vessel dyes to visualize in vivo the vasculature supplying the AP and NTS regions. After identifying and differentiating the mNTS and AP using a combination of approaches, including retrograde neuronal tracing, small-molecule-permeable dyes, and topographical differences in their capillary beds^18,21^, we quantified the functional dynamics of the major arterioles, venules, and capillaries in this region. Previous work has anatomically characterized the microcirculation of the AP and NTS in a mouse model, providing evidence for a continuous microvascular network between these two nuclei^21^. In our in vivo rat model, we confirmed the presence of junction points between the capillary beds of AP and the mNTS previously reported^21^. More importantly, we determined that blood flow between these capillary beds is predominantly unidirectional, from the mNTS to AP. These results support the existence of a functional shared capillary network through which diffusible signals in the NTS can influence the AP.

The vasculature we identify and characterize herein on the dorsal surface of the rat brainstem aligns with that reported in previous anatomical studies in mice using iDISCO tissue-clearing methods^21^. This includes the main parenchymal arterioles and venules that we identified as supplying both the AP and NTS capillary beds. Moreover, we found AP capillaries to be larger than those in the NTS and readily identified junction points between the two capillary beds, supporting a similar neurovascular organization in the mouse and rat brainstem.

An intrinsic limitation of in vivo 2P imaging is that achieving a full depth of field, especially in the NTS, is difficult due to the limits of laser penetration through tissue^30^. Thus, the vasculature supplying this region is not fully mapped in this study. For example, the venule drainage for the NTS capillary bed and other ventral arterioles, previously reported in ex vivo anatomical studies^7,21^, could not be readily visualized in this preparation. Despite these limitations, we are confident that the development of a functional vascular map, as presented here, including the identification of local parenchymal arterioles, which play significant roles in regulating blood supply and flow to a capillary bed, will enable future investigation of neurovascular coupling mechanisms within this region, a phenomenon largely understudied. Given the complex roles both AP and the NTS play in blood pressure sensing and regulation^31^, this region stands as an ideal model for studying the interplay between neurovascular coupling and CNS autoregulatory mechanisms. Thus, future studies addressing this knowledge gap and employing the approach described here are warranted.

The NTS is a heterogeneous nucleus, and with recent advances in genetic tools that enable the generation of cellular atlases, we are beginning to understand the complexity of this neural population^32,33^. In the present study, a specific subset of NTS neurons was labeled: those projecting to the PVN^34^ and SON^35^. We found that many of the retrogradely labeled neurons were catecholaminergic (***Fig.2***), consistent with the substantial A2 noradrenergic projections to both the PVN and SON. These A2 neurons are central to cardiorespiratory and neuroendocrine regulation^36–39^ and receive direct inputs from visceral afferents^40^ and the AP^27^. Not all hypothalamic-projecting NTS neurons are catecholaminergic; a substantial subset expresses other neuropeptides, such as GLP-1^41^, and lies outside the A2 region^27^. We specifically targeted these hypothalamic-projecting neurons because the NTS (but not the AP)^3,27,35^ projects to both the PVN and SON. This strategy allowed for further determination of the anatomical boundary between the AP and mNTS, complementing the clear distinction provided by the AP’s larger, sinusoidal capillaries

Known differences exist between the capillary beds of circumventricular organs and those of classic parenchymal brain capillaries, ranging from structure to function. Specifically, previous reports directly comparing the diameters of capillaries of the AP and NTS in a mouse model reported that the NTS capillaries are significantly smaller than those in the AP^21^. This previous work was performed in perfusion-fixed, empty vessels, a condition in which the lack of flow and tone could impact their measured diameter. Nonetheless, our findings confirm this relationship in vivo (***Fig.4***), in which the vessels are perfused and maintain their tone.

Our observation that capillaries within the AP have diameters nearly twice those in the NTS provides a plausible structural basis for the predominant NTS ➔ AP direction of blood flow we observed at their junctions. Although flow direction is ultimately determined by local pressure gradients, differences in microvascular geometry can strongly influence those gradients by altering the resistance to flow within each network. According to Poiseuille’s law, resistance varies inversely with the fourth power of vessel radius^42^. Therefore, the narrower NTS capillaries impose a substantially greater resistance to flow than the wider AP capillaries. This higher internal resistance likely elevates local upstream pressures within the NTS relative to the AP, particularly at interconnections, favoring blood flow from the NTS toward the lower-resistance AP bed. In this context, the larger, more compliant AP capillaries may act as a *low-impedance sink*, facilitating perfusion transfer from the NTS. The lack of differences in capillary velocities between these capillary beds further supports the notion of the AP acting as a low-impedance sink, capable of accommodating greater volumetric flow without necessitating an increase in capillary velocity. Thus, geometrical differences in capillary diameter may not determine, but likely bias, the directionality of blood flow between these two interconnected microvascular territories.

Prominent neural connections between these two regions have been reported, including a dense innervation from AP neurons to the commissural and mNTS, with comparatively less reciprocal projections from the caudal NTS back to AP^43^. These synaptic connections, including those with the dorsal motor nucleus of the vagus, form the dorsal vagal complex, which plays crucial roles in integrating visceral sensory input with signals from nutrient and hormonal circulation^32^. The recent discovery and characterization of a shared capillary network between the AP and the NTS^21^, including the present work, provides a novel and alternative avenue for inter-region communication and visceral sensory integration. Our work is the first to identify these continuous capillary beds in vivo and to determine the direction of blood flow between them (predominantly NTS ➔ AP). We have previously employed a similar anesthetized rat preparation to define blood flow directionality within the SCN–OVLT vascular portal system^17^ and to quantify changes in neurovascular-coupling dynamics in the SON^23^, and these approaches have yielded reproducible, published results. While anesthesia is known to reduce basal blood pressure and consequently decrease overall cerebral blood-flow rates compared to conscious animals, there is no evidence that it affects the *directionality* of flow within these vascular pathways.

The functional significance of the NTS ➔ AP blood flow remains unknown, but it provides a potential avenue for signals from the mNTS to reach the AP beyond the synapse. The preferential NTS ➔ AP flow suggests that the AP, despite being a circumventricular organ exposed to circulating signals, may also receive significant metabolic and paracrine influences from the NTS microvasculature. Such a configuration could facilitate the transfer of diffusible signaling molecules from the NTS into the AP without dilution in the systemic circulation. Conversely, the low-resistance nature of the AP vasculature could facilitate pressure dissipation and venous drainage in the NTS, thereby maintaining stable perfusion in this critical integrative center. Together, these findings reveal an unappreciated level of vascular specialization within the dorsal medulla, in which differences in capillary geometry could contribute to shaping local hemodynamics and directional, flow-dependent communication between the NTS and AP.

Lastly, it is important to consider that capillary blood flow direction is not always constant, as flow within an individual capillary can stall and even reverse in pathological conditions^44^. Thus, it would be important to assess whether blood flow and directionality within the AP-NTS shared capillary network are altered in disease states involving the dorsal vagal complex, such as hypertension, heart failure, and obesity, among others.

## Acknowledgements

This work was supported by NHLBI 162575 (JES) and by the Office of The Director, National Institutes Of Health and National Institute Of General Medical Sciences under Award Number S10OD032336-01.

---

**Supplementary Figure 1:**
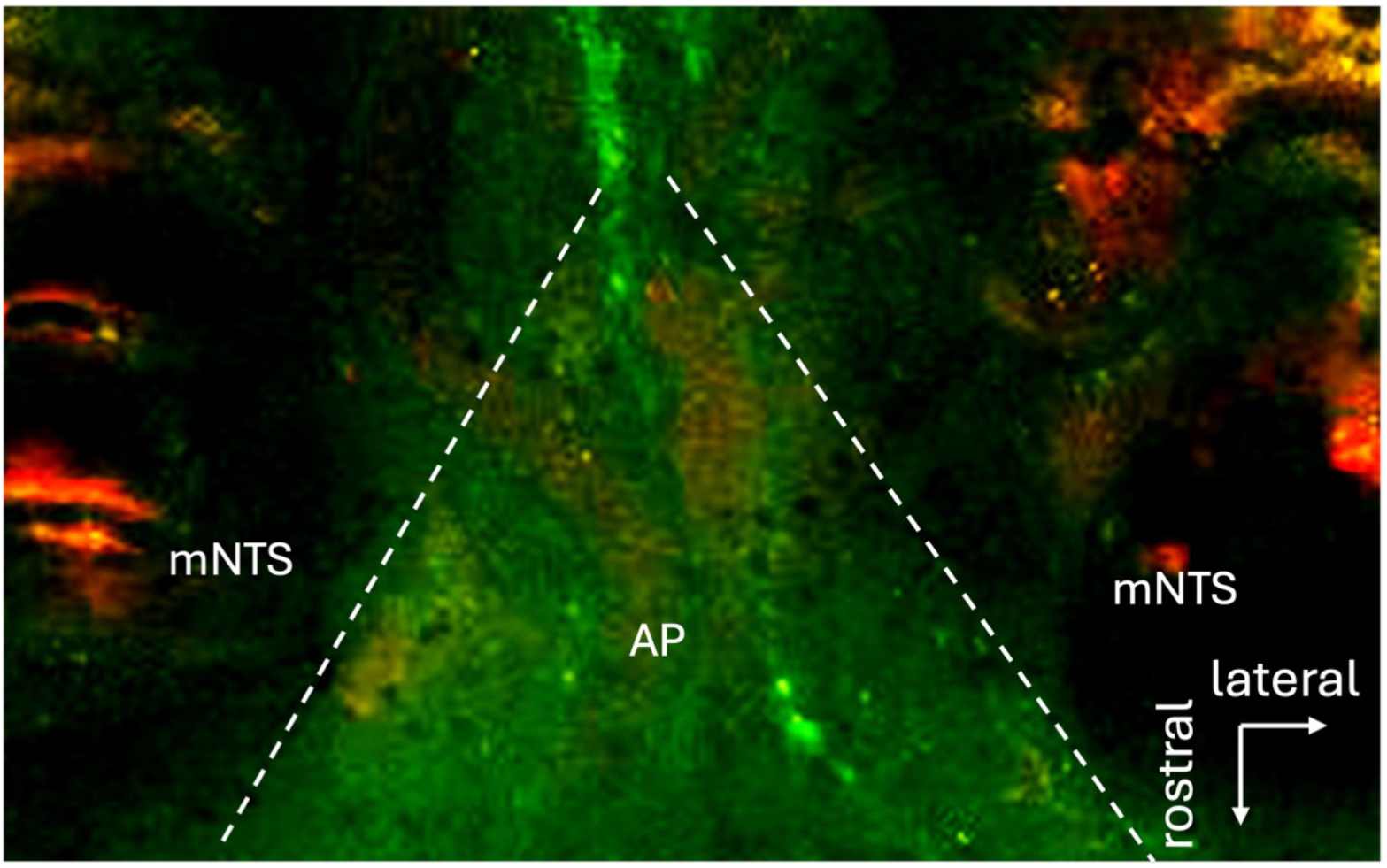
NaF further defines the border between AP and the mNTS. Representative image of the dorsal brainstem after intravenous infusion of sodium fluorescein (NaF, green) and rhodamine (red). NaF is seen excised from the vessels, as indicated by the increased density of green fluorescence, found near the midline, where the enlarged capillaries of AP reside. Dashed white lines indicate the borders of AP as delineated by the NaF.

## Notes

### Competing Interest Statement

The authors have declared no competing interest.

